# Evaluation of the potential food allergy risks of recombinant human lactoferrin expressed in *Komagataella phaffii*

**DOI:** 10.1101/2024.01.18.576250

**Authors:** Yanisa Anaya, Raysa Rosario Martinez, Richard E. Goodman, Philip Johnson, Shashwat Vajpeyi, Xiaoning Lu, Ross Peterson, Sarah M. Weyers, Bella Breen, Kahler Newsham, Brian Scottoline, Anthony Clark, Carrie-Anne Malinczak

## Abstract

Prior to the introduction of novel food ingredients into the food supply, safety risk assessments are required, and numerous prediction models have been developed and validated to evaluate safety. The allergenic risk potential of Helaina recombinant human lactoferrin (rhLF, Effera™), produced in *Komagataella phaffii* (*K. phaffii*) was assessed by literature search, bioinformatics sequence comparisons to known allergens, glycan allergenicity assessment, and a simulated pepsin digestion model. The literature search identified no allergenic risk for Helaina rhLF, *K. phaffii*, or its glycans. Bioinformatics search strategies showed no significant risk for cross-reactivity or allergenicity between rhLF or the 36 residual host proteins and known human allergens. Helaina rhLF was also rapidly digested in simulated gastric fluid and its digestibility profile was comparable to human milk lactoferrin (hmLF), further demonstrating a low allergenic risk and similarity to the hmLF protein. Collectively, these results demonstrate a low allergenic risk potential of Helaina rhLF and do not indicate the need for further clinical testing or serum IgE binding to evaluate Helaina rhLF for risk of food allergy prior to introduction into the food supply.

## 1. INTRODUCTION

Lactoferrin (LF) is an 80 kDa, iron-binding glycoprotein found in exocrine secretions of humans and many other mammals, predominantly milk and colostrum in humans, and in colostrum in ruminants. The roles of this bioactive protein are multifunctional, including linking innate and adaptive immune responses, iron homeostasis, and many other functional benefits.^1^ Lactoferrin is produced in the epithelial cells of mammary, salivary and lacrimal glands as well as in neutrophils. Composed of 691 amino acids, human milk lactoferrin (hmLF) plays several critical roles, including but not limited to, the neonatal immune system and iron homeostasis.^1–3^ Bovine lactoferrin (bLF) is a 689 amino acid polypeptide sharing 69% sequence homology with hmLF, and shares a number of beneficial properties with hmLF. Bovine LF is generally recognized as safe (GRAS) (US FDA, GRN 000067, 2001; US FDA, GRN 000077, 2001c; US FDA, GRN 130, 2003; US FDA, GRN 000423, 2012; US FDA, GRN 000464, 2013a; US FDA, GRN 000465, 2013b; US FDA, GRN 000669, 2016)^4–10^ for use in cow’s milk-based term infant formula, sports and functional foods, ice cream, powdered milk, yogurt chewing gum, and as an antimicrobial agent.^11^ In addition to the intended uses described above, bLF is used in cosmetics and dietary supplements. While bLF is effective as a dietary ingredient, its bioavailability and subsequent functions may differ from human LF (hLF). For example, in vitro studies have shown that hLF digests more slowly and has a stronger affinity for the human LF receptor in the small intestine, indicating that it may be more bioavailable than bLF.^12–14^ Thus, utilizing hLF as a substitute for bLF could be more appropriate. However, isolating LF from human milk is impractical for this purpose because it is found in relatively low concentrations, resulting in high production costs as well as the ethical challenges of using human milk to isolate LF rather than to feed infants. Strategies to produce cost-effective, sustainable, and safe hLF are warranted to develop successful commercial production.

Recombinant human lactoferrin (rhLF) has been produced in several biological expression systems, including, but not limited to, rice,^15^ transgenic cows,^16^ Chinese hamster ovary (CHO) cells,^17^ and yeast, namely *Komagataella phaffii (K. phaffii)* (previously known as *Pichia pastoris*).^18^ Introduced as a commercial biological expression system in the 1980s,^19^ *K. phaffii* is a methylotrophic yeast that remains one of the most popular fermentation systems for recombinant protein synthesis in molecular biology due to its ease of use, standardized protocols, scalability and capabilities to appropriately fold and secrete desired proteins.^20^ Food ingredients available for specific intended uses in the US food supply that use *K. phaffii* for protein production include β-lactoglobulin^21^ (Remilk), egg white protein^22^ (The EVERY company), myoglobin^23^ (Motif FoodWorks) and soy leghemoglobin^24^ (Impossible Foods). Furthermore, *K. phaffii* is an approved protein source for broiler chickens at up to 10% of the total feed.^25^

Helaina Inc. (New York City, New York, USA) successfully developed rhLF derived from a glycoengineered yeast that is substantively similar in structure to hmLF.^26^ Helaina rhLF (Effera™) was engineered according to the UniProt protein ID for human LF, TRFL_HUMAN (P02788). A proprietary technology, involving disruption of an endogenous glycosyltransferase gene (OCH1)^27^ and a stepwise introduction of heterologous glycosylation enzymes, enables a modified strain of *K. phaffii* (GS-115) to produce rhLF.^28^ It has been well established that hmLF is a glycoprotein mainly having complex N-linked glycans on three sites including Asn-156, Asn-497 and Asn-642.^29^ Helaina rhLF possesses glycans on the exact same three sites with primarily oligomannose N-glycans having 5 to 14 mannose residues. A thorough comparison of Helaina rhLF glycosylation profile compared to hmLF is described in detail elsewhere.^26^

Evaluating the food allergy risk of novel proteins prior to their introduction into the food supply is a best practice. While there is no definitive, single factor that serves as an indicator of protein safety, the Codex Alimentarius Commission’s multistep approach provides a framework by which to evaluate potential allergenic risk and the safety profile of novel food proteins in genetically modified microorganisms.^30^ However, it is important to recognize that Codex was not intended to evaluate novel foods that can have tens to hundreds of new proteins that may be evolutionarily conserved in multiple taxa, and often of low abundance in food products. Codex guidelines recommend reviewing the safety of the protein gene source and history of exposure, comparing the amino acid sequence similarity of the novel protein to known human allergens, and testing the stability of the protein to digestion by pepsin to provide an overall assessment on allergenic potential. Although other methods (e.g., animal models and targeted sera screening) are being developed to test for food allergy, such approaches have not been thoroughly evaluated or validated for predicting protein allergenicity.^31–33^

Codex guidelines have been applied to evaluate the potential allergenicity of other novel proteins expressed in a recombinant host and residual host proteins that remain in food fractions, such as rhLF produced in transgenic cows^34^ and soybean leghemoglobin.^35^ However, the guidelines have yet to be applied to rhLF expressed in *K. phaffii*; therefore, the main objective of this work was to apply Codex criteria to evaluate the potential allergenic risk of Helaina rhLF. Additionally, an in vitro pepsin digestion model developed by Astwood et al.^36^ and refined by Thomas et al.,^37^ and Ofori-Anti et al,^38^ which is widely used to simulate the digestion of proteins to give an indication of risk predictions (e.g., allergenicity), was performed. Although not part of the Codex weight-of-evidence approach, allergenicity based on the glycan signature of Helaina rhLF was also assessed due to the glycoprotein nature of hLF.

## 2. MATERIALS AND METHODS

### 2.1 Scientific literature review - Evidence of allergenicity

The National Center for Biotechnology Information (NCBI) PubMed® literature database was searched on October 26, 2023, for peer-reviewed evidence of allergenicity to hLF as well as proteins from *K. phaffii* and from its previous name, *Pichia pastoris,* using keyword limits. The searches used combinations of species names “*Komagataella phaffii*” and “*Pichia pastoris*” with either “clinical allergy” or “food allergy” or “IgE binding”.

The potential for cross reactivity against specific glycans introduced by the yeast expression system (i.e., mannose) was also assessed to determine if a risk of allergenicity against asparagine-linked glycans exists. This risk includes identifying the presence or absence of cross-reactive carbohydrate determinants (CCD) that are protein-linked carbohydrate structures (i.e., glycans) involved in the phenomenon of cross-reactivity of sera from allergic patients towards a wide range of allergens typically found in plants and insects.

### 2.2 Allergenicity assessment of relatively abundant residual proteins from Komagataella phaffii

#### 2.2.1 Sequence of rhLF and residual K. phaffii host proteins

Three representative production batches of Helaina rhLF (UniProt P02788) were analyzed by liquid chromatography-tandem mass spectrometry (LC-MS/MS). Three replicate analyses of each production batch were performed.

The LF samples were solubilized in a 50 mL Tris HCl buffer (pH 8.0). Samples were diluted in 25 mM ammonium carbonate and 10 mM dithiothreitol (DTT) at 60 °C, followed by addition of 50 mM iodoacetamide (IAA) at room temperature. Diluted samples (10 μg) were digested with trypsin enzyme (Promega, Madison, WI, USA) at 37 °C for 18 hours, quenched with formic acid, and desalted using SPE.

Peptide digests were analyzed by nano-flow LC-MS/MS with a Waters M-class high pressure liquid chromatography (HPLC) system interfaced to an Orbitrap Fusion Lumos mass spectrometer (Thermo Fisher Scientific, Waltham, MA, USA). Peptides were loaded onto a trapping column and eluted over a 75 μm analytical column at 350 nL/min; both columns were packed with Luna C18 resin (Phenomenex, Torrance, CA, USA). A 4-hour gradient was applied. The mass spectrometer was operated in data-dependent mode, with MS and MS/MS performed at resolutions of 60 000 full-width at half maximum (FWHM) and 15 000 FWHM, respectively. Advanced peak determination was enabled. The instrument was run using a 3-second cycle for MS and MS/MS. Only the peptides having 2 to 7 charges were selected for MS/MS and were excluded for 25 s from repeated MS/MS. The fragments were generated by higher-energy collisional dissociation (HCD) with a fixed normalized collision energy at 30. Orbitrap was used as the detector with a resolution of 15,000, maximum injection time 22 millisecond, and Automatic Gain Control target 50000, respectively.

Data-dependent acquisition mass-spectral data was analyzed using PEAKS version 8.5.^39^ The sequence dataset used for the analysis was the publicly available *K. phaffii* set from UniProt (Taxon ID 4922, containing 5,257 entries) appended with the amino acid sequence of hmLF (UniProt P02788) and mannosidase from *Trichoderma reesei* (*T. reesei*) (UniProt G0RBB5). The following data analysis parameters were used: no mis-cleavages, carbamidomethylation as a fixed modification, oxidation of methionine as a variable modification, parent mass error tolerance of 5 ppm, fragment mass error tolerance of 0.02 Da, false discovery rate (FDR) of 1% and charge states between 2 and 4 were accepted. Only proteins identified via two or more peptides were considered for further analysis. The resultant lists of protein IDs (one for each replicate analysis for each batch) were combined for use in comparison to allergenic proteins in the www.AllergenOnline.org version 22 database, as described below.

After protein identification, label-free quantitation was performed using peptide ion (MS1) abundance to allow for the determination of the quantity of *K. phaffii* proteins relative to the quantity of rhLF, and determination of which residual *K. phaffii* proteins from strain GS-115 were most abundant. Abundance data was not used to select proteins for comparison to allergenic proteins.

#### 2.2.2 AllergenOnline, version 22

The www.AllergenOnline.org database (version 22, released on May 25, 2023) is a publicly available tool that provides a peer-reviewed allergen list and sequence database to identify proteins that may pose risk of allergenic cross-reactivity. Updated annually, the website is designed to help assess the safety of novel proteins developed through genetic engineering or food processing prior to their introduction into the food supply.^40^

Identified proteins from UniProt that were matched by LC-MS/MS were queried against allergens in AllergenOnline to determine significant alignment with known allergens. The full amino acid sequence of human lactoferrin and each identified *K. phaffii* protein (strain GS-115) and *T. reesei* mannosidase were entered into the sequence search window of AllergenOnline using the full-length FASTA, version 36 search. A separate search was performed for 80 amino acid alignments using FASTA, which uses a sliding window of 80 amino acid segments of each protein to find identities of greater than 35%. Matches to allergenic proteins were evaluated further using publications in the AllergenOnline.org database associated with the matched allergens, as well as consideration of the breadths of taxonomic matches to the same protein by BLASTP.

#### 2.2.3 BLASTP in NCBI Entrez Protein Database

The Basic Local Alignment Search Tool (BLAST®) is a publicly available tool on the NCBI website (http://www.ncbi.nlm.nih.gov/BLAST/) that identifies regions of local similarity between gene or protein sequences. Protein Blast (BLASTP) identifies a protein sequence against the NCBI Entrez Protein Database, a search and retrieval system that allows users to access information from many health science databases to provide information on protein structure, function, and evolution.

Given the frequency by which BLASTP is updated compared to AllergenOnline (daily or weekly, versus annually), this tool was utilized to identify newly discovered allergens. The identified proteins were queried in BLASTP, version 2.14.01, on September 26, 2023, to check for allergens that may have been missed in AllergenOnline. Two searches were conducted; first, without any keyword limit looking for the best identities, and second, with a keyword limit of “Homo sapiens species taxonomic ID 9606”, to consider likely tolerance limits.^41^ The primary criterion to judge the significance for any alignment was whether the BLASTP result for “no keyword” was higher than BLASTP of the protein to human proteins. In addition, high identity matches across diverse taxa are highly unlikely to act as clinically relevant allergens based on restricted clinical mandates from the allergy literature.

### 2.3 In Vitro Pepsin Digestibility Study

A simulated in vitro digestion study was performed to determine the stability of Helaina rhLF in simulated gastric conditions, compared to hmLF and bLF, in the presence of pepsin enzyme per Codex guidelines. Experimental procedures were modeled after those developed by Astwood et al.,^36^ and refined by Thomas et al.,^37^ and Ofori-Anti et al.^38^

#### 2.3.1 Preparation of LF (rhLF, hmLF, and bLF), simulated gastric fluid, and controls

Recombinant human lactoferrin (Helaina rhLF), α-isoform, was produced by fermentation and expressed from a *K. phaffii* (GS-115) recombinant system, purified by microfiltration/diafiltration and cation exchange chromatography, and then spray-dried to a powder. The purity was measured by high performance liquid chromatography (HPLC) and sodium dodecyl sulfate–polyacrylamide gel electrophoresis (SDS-PAGE) and is typically greater than 95%. The iron saturation was determined to be 58% saturated.^26^ Helaina rhLF was reconstituted and diluted into phosphate buffered saline (PBS) prior to use.

Lactoferrin was isolated from human milk that was prepared by the Northwest Mothers Milk Bank and provided by Brian Scottoline, MD PhD at Oregon Health & Science University (OHSU). The milk from six donors was pooled and frozen prior to use. The hmLF was then isolated from the milk using the following procedure: milk samples were centrifuged at 10,000xg for 30 minutes and the liquid fraction removed. The resultant liquid fraction was adjusted to pH 4.7 before incubation at 40℃ for 30 minutes for casein precipitation. The solution was then centrifuged at 10,000xg for 30 minutes at 4℃ to separate liquid and solid fractions. The supernatant was decanted and purified via ultrafiltration/diafiltration (UFDF) and chromatography as described above for rhLF. The purified product was then buffer exchanged into 1X PBS pH 7.4 (no spray drying) and concentrated against a 30kDa membrane. This procedure yielded approximately one gram of LF per liter of human milk with a purity of greater than 95% as measured by HPLC and SDS-PAGE. The measured iron saturation was ∼30%. This work was not determined to be human research by the OHSU Institutional Review Board.

Bovine lactoferrin (bLF) isolated from bovine milk was purchased from Lactoferrin Co, Australia (Product 11683) and reconstituted into PBS prior to use. The purity provided by the supplier was greater than 95% and iron saturation was 9.9%.

Lyophilized β-lactoglobulin (Sigma, #L3908) was reconstituted with PBS and the concentration (mg/mL) was verified via Nanodrop® (Model: NanoDrop™ One/OneC, Thermo Fisher Scientific).

Simulated gastric fluid (SGF) was comprised of 0.084 N HCl, 35 mM NaCl, pH 2.0, and 3200 units (U) porcine pepsin (Sigma, P6887, 4048 U/mg activity) to obtain 10 U pepsin activity per µg protein.

The protein positive (no pepsin) and negative (no protein) controls were SGF containing 0.2 mg/mL LF (or β-lactoglobulin) without pepsin, and SGF containing 3200 U of pepsin with no LF, respectively. Positive controls were prepared for each experimental sample.

#### 2.3.2 Experimental Procedures

Each experimental tube contained a total volume of 1.6 mL of SGF, 0.2 mg/mL of LF (rhLF, hmLF, or bLF), and 3200 U of pepsin to create a ratio of 10 units of pepsin to 1 µg protein. β-lactoglobulin (0.2 mg/mL) was used as a non-digestible control. Pepsin was added immediately to each tube and mixed. Tubes were incubated at 37°C and 100 μL aliquots were removed after specified sample times: 0.5, 2, 5, 10, 20, 30 & 60 minutes. Pepsin activity was quenched using 35 uL of 1M NaHCO_3_, pH 11 for every 100 μL of the experimental digestion aliquot. For the 0 time-point control, pepsin activity was quenched prior to the addition of protein.

Samples from each digestion time-point, including all controls, were run on a tris-glycine gel to visualize the degradation of the rhLF in comparison to hmLF and bLF. Additionally, all proteins were visually compared to a corresponding 10% protein control (0.02 mg/mL) to ensure a ≥ 90% digestion profile of each protein. To determine the detection limit, a serial dilution of protein samples was prepared with PBS covering the range of 200% total protein (0.4 mg/mL) per well, to 5% (0.01 mg/mL) per well. Samples were separated in an SDS-PAGE at a constant 185V for 45 min, followed by staining with Imperial Protein Stain.

#### 2.3.3 Automated Western Blot

The Jess™ Simple Western System (ProteinSimple, San Jose, CA, USA) is an automated protein separation and immunodetection system. To visualize and quantify the digestion of proteins in SGF, manufacturer’s instructions for 12-230 kDa Jess™ separation module [Ref-No. SM-FL004] were followed. The experimental samples were diluted to 0.004 mg/mL with distilled water, and then 1 part of 5x Fluorescent Master Mix was combined with 4 parts of the diluted sample to achieve a final concentration of 0.0032 mg/mL. Samples were denatured for 10 min at 90 °C and centrifuged for 5 minutes at 2500 rpm (∼1000 x g). The fluorescent molecular weight marker, (12-230 kDa [Biotinylated Ladder]), and sample proteins were separated in capillaries as they moved through a separation matrix at 375V. Jess™ immobilized the protein samples to the capillary walls using UV light. In addition, it automatically incubated the samples in blocking solution (Antibody Diluent 2, ProteinSimple, Cat-No. 042-203), primary antibody (anti-human lactoferrin antibody, Sigma, Cat-No. L3262), secondary antibody (Goat-Anti-Rabbit Secondary Antibody, Protein Simple, Cat-No. 040-656), and performed the necessary washing steps between incubations. Chemiluminescence was established with Luminol-Peroxide Mix (Luminol-S, ProteinSimple, Cat-No. 043-31; Peroxide, ProteinSimple, Cat-No. 043-379). Digital imaging was captured with Compass Simple Western™ software (version 6.2.0, ProteinSimple) to produce gel images and histograms. Relative quantification was measured by calculating the percent (%) area of the chromatogram peaks.

#### 2.3.4 Conventional Western Blot

Samples were separated in an SDS-PAGE at a constant 185V for 45 min and the gels were removed from the cassette and the iBlot 2 Gel Transfer was used to transfer the proteins to polyvinylidene difluoride (PVDF) membranes. The membranes were placed in an opaque gel box and incubated with blocking buffer (PBST + 10% Fish gelatin (PBS 10X, pH 7.2, Thermo Fisher Scientific, Cat-No. 70-013-032; Biotium 10X Fish Gelatin Blocking Agent, Thermo Fisher Scientific, Cat-No. NC0382999) at room temperature for 1 hour. Samples were washed with PBST (PBS + 0.1% Tween 20 [Tween 20, Sigma-Aldrich, Cat-No. P9416-100ML]) and then incubated with the primary antibody (rabbit anti-human lactoferrin antibody; Sigma, Cat-No. L3262) for 1 hour at room temperature. Samples were washed and incubated with the goat anti-rabbit Alexa-Fluor secondary antibody (Thermo Fisher Scientific, Cat-No. A-11034) at room temperature for 30 minutes. Samples were washed 5 times, then rinsed for 2 minutes in distilled water to stop the reaction. Results were visualized using the iBright imager (Thermo Fisher Scientific) and the settings: Alexa Fluor 488, No false colors, exposure time: 00:00 min: sec.

## 3. RESULTS

### 3.1 Literature Searches Identify No Allergenic Risk for rhLF, K. phaffii, or its Glycans

Literature searches in PubMed for the primary protein of interest, hLF, did not identify studies showing allergy to this protein, even though the amino acid sequence is moderately identical to bovine lactoferrin (70% full-length identity) and ovo-transferrin (52% overall identity), known potential allergens. The literature results included a study on rhLF from transgenic cows that underwent an allergenicity assessment using the Codex guidelines and included a bioinformatics analysis, stability testing of rhLF in pepsin, and serum reactivity tests to evaluate potential allergenicity.^34^ Considering the rapid digestion by pepsin and no specific binding of IgE using serum from patients with milk and egg allergy, the authors concluded the allergenicity potential of this self-protein is very low.^34^ A single paper that suggested a possible risk from human lactoferrin was for protein expressed in rice that has cross-reactive carbohydrate determinants (CCD) on asparagine-linked glycosylation sites which may be linked to allergy.^42^ Importantly, the rice-expressed protein did not elicit basophil activity due to IgE binding to CCD and there was no proof of allergy to that product. In addition, the IgE binding described in the paper on rice was to complex carbohydrates with α(1-3)-fucose and/or β(1-2)-xylose on the stem loop, which Helaina rhLF does not contain.

Literature searches were also conducted for peer-reviewed evidence of allergenic risk to proteins from *K. phaffii* and *P. pastoris.* The searches used combinations of species names “*Komagataella phaffii*” and “*Pichia pastoris*” with either “clinical allergy” or “food allergy” or “IgE binding”. Keyword limits “allergy” and “*K. phaffii*” identified one publication.^42^ “*P. pastoris”* identified 156 publications, including Jin et al. on production of soybean leghemoglobin in the same yeast expression system.^35^ The other 155 publications describe cloning and expression of various plant and fungal proteins for laboratory testing including major allergens such as the 2S albumin Ara h 6 from peanuts. Using the full Latin name “*Komagataella phaffii*” and “clinical allergy” did not identify any papers. Using the search terms “*Pichia pastoris*” and “clinical allergy” identified 34 papers including one with direct administration of recombinant human serum albumin made in this yeast, injected three times over 3 days, in 423 cirrhosis patients with ascites or edema.^43^ There were 96 adverse reactions from the treatment but no allergic reactions or IgE binding to yeast proteins reported. Furthermore, the injection administration route of recombinant human serum albumin is not applicable to Helaina rhLF that would be taken orally as a dietary protein and digested through the gastrointestinal tract.

Information relative to the expression system, *K. phaffii*, was considered, including potential addition of glycans. While *K. phaffi* produces oligomannose glycans that can be bound by mannose receptors on dendritic cells (DC), Kreer, et al. found no proof for the DC triggering immunogenic and/or subsequent allergic reactions.^44^ Glycans that have been implicated in IgE binding are CCD that are common on many plant-produced glycoproteins or α(1,3)-galactose (α-gal) sugars added to proteins in tick saliva and on some red meat proteins.^42,45^ Importantly, Helaina rhLF does not contain any of these CCD and the literature does not support allergenicity of the oligomannose N-glycans that make up Helaina rhLF.^26^

### 3.2 Bioinformatics search strategies show no significant risk for cross-reactivity and allergenicity

Protein identification on three individual batches of Helaina rhLF was performed using LC-MS/MS. This analysis confirmed the expected sequence for rhLF with full protein coverage excepting signal sequence and those peptides unlikely to be detected due to size of peptides (m/z cutoff). Label-free quantitation indicated that ≥99% of the identifiable protein content was, in fact, the target protein, human lactoferrin (Table 1, Figure 1A, Supplemental Data). Comparison of hLF sequence to AllergenOnline identified protein matches of 70% identity to *Bos taurus* (bovine) lactotransferrin (lactoferrin), which is a minor milk allergen, and 52% identity to ovo-transferrin, a minor food allergen in chicken eggs by overall FASTA (Table 2). However, by sliding 80 AA window the identities were higher; bovine lactotransferrin was 83.8% with rhLF and there were 631 matched 80mers compared to 69.7% overall identity to rhLF. The best 80mer match was to ovo-transferrin at 67.6% ID with 631 matches of 80 AA, compared to 51.8% overall ID by FASTA. The remaining <1% protein content included 36 host cell proteins (Table 1), with no single protein having an average relative abundance greater than 0.05%. Some variability in host cell proteins was observed between the three batches (Figure 1B); however, no single protein had greater than 0.075% relative abundance in any single batch (Table 1).

**Figure 1.**
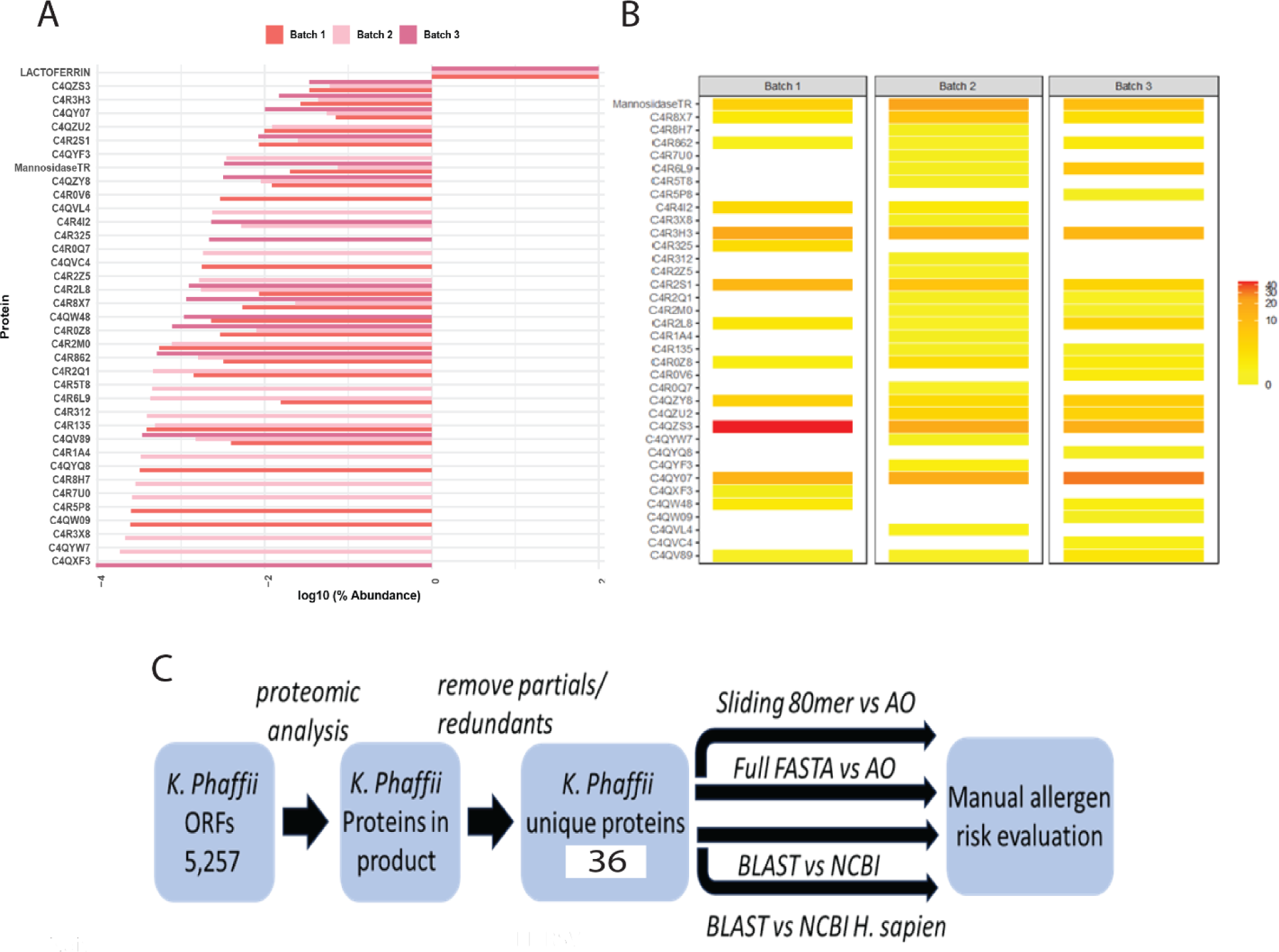
Bioinformatics search strategies show no significant risk for cross-reactivity and allergenicity. **A.** Relative abundance of rhLF compared to K. Phaffii host proteins on a log 10 scale in three representative batches. Lactoferrin comprises more than 99% of the total proteins in these batches**. B.** Heatmap showing abundance of host cell proteins**. C.** Workflow to evaluate allergenic risk of residual *K. phaffii* proteins. Mass spectrometry was used to determine the presence of proteins in rhLF product from a database of *K. phaffii* ORFs. After removing redundant sequences and proteins not specific to *K. phaffii* GS-115 or *T. reesei* (mannosidase), 36 protein sequences were used to query AllergenOnline (AO) using identity in a sliding 80mer search and by full-length FASTA identity search. The 36 protein sequences were also used in a BLASTP search of NCBI proteins, and in a separate NCBI search limited to *H. sapiens* proteins. Evaluation of risk is manual and is informed by the results of these four sequence identity comparisons.

**Table 1.**
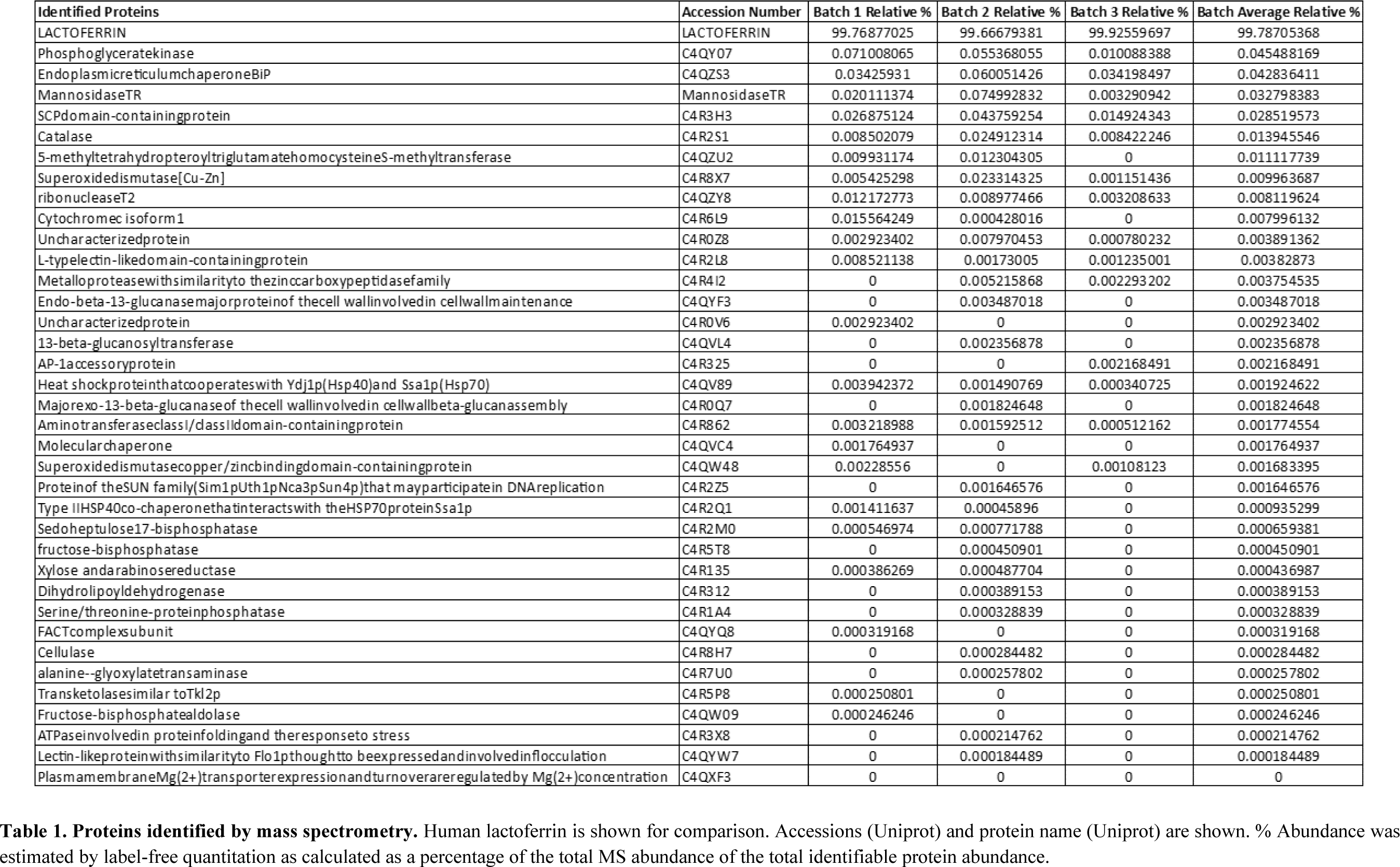
Proteins in Helaina rhLF Product.

**Table 2.** Recombinant Human Lactoferrin Allergen Assessment.

| Accession number | FASTA of NCBI Protein | 80mer Match | Best overall FASTA vs v22 | BLASTP vs Homo sapien | Is there RISK of allergy? |
| --- | --- | --- | --- | --- | --- |
| P02788 | Lactoferrin | 83.8% ID to *Bos Taurus* lactotransferrin | 70% ID to *Bos Taurus* lactotransferrin | 100% ID human lactoferrin | **NO tolerance to human** |
|  |  | 67.6% ID to *Bos Taurus* lactotransferrin | 52% ID to *Bos Taurus* lactotransferrin | 100% ID human lactoferrin | **NO tolerance to human** |

A bioinformatics search strategy was used to determine amino acid sequence homology and structural similarity of the identified *K. phaffii* proteins (strain GS-115) to known human allergens. The amino acid sequences of 36 unique proteins were identified by LC-MS/MS (Table 1, Supplemental Data) and were verified by UniProt accession numbers and searched using AllergenOnline and BLASTP. The workflow for protein identification and subsequent risk evaluation by homology searching is shown in Figure 1C. The overall results of the bioinformatics comparisons to AllergenOnline by FASTA and to NCBI Protein database by BLASTP are defined in Supplemental Table 1. Importantly, of these 36 identified proteins, none were linked with a significant risk for cross-reactivity and allergenicity.

The most abundant host cell protein identified was phosphoglycerate kinase at 0.045% of the protein content (Table 1) and no allergen match by sliding 80 AA window (Table 3). It had a best overall FASTA alignment to α-s1 casein at 23% overall identity. The BLASTP was to itself and the best human alignment was 65% ID to human phosphoglycerate kinase. Proteins with high identity to human proteins are likely to be tolerated and this identity match is much higher than to the milk allergen α-s1 casein and presents low risk of allergic cross-reactivity. The second most abundant host cell protein was endoplasmic reticulum chaperone BiP protein (Table 1), found to have 84%-36% ID to heat shock protein (HSP) of mosquito, mites, fungi, and grass with BLASTP to itself and 65.6% ID to human HSP, indicating likely tolerance due to close identity to human (Table 3). Mannosidase from *T. reesei*, which is a gene added to *K. phaffii* to limit the potential for hypermannosylation^46^ was the next most abundant protein (Table 1) and was determined to be of low allergenic risk due to low abundance (0.033%), no match by sliding 80 AA window, and 31% ID to the human homologue (Table 3). Of the remaining host cell proteins (Table 1), five were found to have >50% 80mer match to a known or putative allergen (Table 3, Supplemental Table 1). Catalase (0.014%) was found to have a match to a potential allergen, penicillin citrinum catalase, at 60% identity; however, this protein was at low abundance of 0.03% and had high identity (51%) to human catalase, indicating likely tolerance. 5-methyltetrahydropteroyltriglutamate-homocysteineS-methyltransferase which showed 77.5% ID to Sal k 3 pollen allergen was also identified, with 48% ID overall FASTA to that protein and 100% ID to *K. phaffii* but no matches to human protein. By BLASTP this protein had 88% ID to 100 enzymes of a wide variety of plants, trees, and weeds; there is only one laboratory that has published data on IgE binding to this pollen protein as an allergen.^47^ It is highly unlikely there is any clinically identifiable co-reactivity as it is a low abundance enzyme (<0.011%), therefore limited risk. Cytosolic superoxide dismutase was also identified (0.010%) with 62% identity to ambrosia and olive superoxide dismutase and 100% identity to *K. phaffii* superoxide dismutase. However, this protein also had 59% identity to human superoxide dismutase suggesting tolerance due to human recognition. Cytochrome c isoform 1 (0.008%) with one 80mer match of 74% ID to *Curvularia lunata* cytochrome c, full ID to *Curvularia lunata* cytochrome c, and 100% ID to *K. phaffii* and 65% ID to human cytochrome c as well as many other proteins was also present. An ATPase involved in protein folding (0.0002%) was also identified with 91% identity to *Davidiella* HSP and 14 other proteins at lower ID, 100% identity to *K. phaffii* ATPase with 75% identity to human HSP70 suggesting tolerance due to human recognition. The remaining host cell proteins identified (Table 1) had <50% 80mer match to any allergen and therefore, have low allergenic risk (Supplemental Table 1).

**Table 3.** Allergenic assessment of select host cell proteins.

| Accession number | FASTA of NCBI Protein | Relative Abundance | 80mer Match | Best overall FASTA vs v22 | BLASTP vs Protein | BLASTP vs Homo sapien | Is there RISK of allergy? |
| --- | --- | --- | --- | --- | --- | --- | --- |
| C4QY07 | Phosphoglycerate kinase | 0.045% | No Match | alpha s1 casein 23% ID | >CCA37205.1 3-phosphoglycerate kinase [Komagataella phaffii] | 65% ID human Phosphoglycerate kinase | **NO; <50% ID and high ID to human protein** |
| C4QZS3 | Endoplasmic reticulum chaperone BiP | 0.043% | 84% down to 36% to HSP of mosquitoes, mites, fungi, grass | 65% down to 46% to HSP of mosquito, mites, fungi, grass | 100% Komagataella p and down to 75% other fungi | 65.6% HSP | **NO tolerance to human** |
| GORBB5 | Mannosidase (*T. reesei*) | 0.033% | No matches | No matches by FASTA | 100% to Trichoderma reesei full length down to 58% id to other fungal homologous proteins | 31% ID to human protein | **NO tolerance to human** |
| C4R2S1 | Catalase | 0.014% | 60% ID to Penicillium citrinum catalase over 80 AA and 37% ID of that protein by Full-FASTA | 37% ID penicillium catalase and less for 3 species | 100% to Komagataella p down to 65% ID for 100 fungal proteins | 51% down to 41% to 9 human proteins catalase | **NO tolerance to human** |
| C4QZU2 | 5-methyltetrahydropteroyltriglutamate-homocysteine S-methyltransferase | 0.011% | 77.5% ID to same two proteins as by overall FASTAS, Sal k 3 and other | 49% ID to Sal k 3 and 48% to another methyltetrahydropteroyltrig protein | 100% ID Komagataella cobalamin down to 77% ID to many fungal proteins | No Match | **NO; Low Abundance** |
| C4R8X7 | Cytosolic superoxide dismutase | 0.010% | 62% to Ambrosia Superoxide dismutase and to 57% to 23 Olive SOs | 59% ID to Ambrosia So dismutase and 52% or higher to olive | 100% ID to Komagataella p down to 75% ID 100 fungi | 59% ID to human Superoxide dismutase down to 44% ID to other human Sos | **NO tolerance to human** |
| C4R6L9 | Cytochrome c isoform 1 | 0.008% | 74% ID to Curvularia cytochrom c | 71% ID to Curularia lunata cytochrom c | 100% ID to Komagataella cytochrome c and 80% ID to many other fungal cytochromes | 65 id TO HUMAN cytochrome c multiple proteins (genes) and heat shock protein with more than 55, 14, >52% ID | **NO tolerance to human** |
| C4R3X8 | ATPase involved in protein folding | 0.0002% | 91% ID to Davidiella HSP and 14 more to lower identities | Heat shock proteins Tyrophagus 76% ID on down to 46% ID pollen HSP | 100% ID Komagataella ATPase and HSP 85% ascoidea down different fungi ~ 100 with down to 83% identity fungal proteins | 75% ID to heat shock protein 70 Homosapiens and 100 to 45% ID | **NO tolerance to human** |

### Helaina rhLF Digests in a Comparable Manner to Human Milk LF and Shows Low Allergenic Risk Potential in the Pepsin Digestion Model

Previous studies have shown that pepsin digestion of proteins in a simulated gastric fluid (pH 2.0) can be used to correlate allergenic risk potential.^48,49^ In order to assess the digestibility of Helaina rhLF, an in vitro digestion study was performed in SGF, pH 2.0 with a pepsin:protein ratio of 10 units pepsin per 1 µg of protein. Successful digestion to ≤10% of the initial starting concentration indicates low allergenic risk potential.^38^ Initially, to evaluate staining characteristics of each test protein and to ensure detection down to 10% of the target protein, a preliminary SDS-PAGE-stained gel with samples of serially diluted target protein was performed and showed similar detection levels of all proteins tested (Supplemental Figure 1 A, B). Next, the pepsin digestion study was performed as previously described to test digestibility of rhLF compared to hmLF and bLF. These results indicate that Helaina rhLF digests in a comparable manner to LF isolated from human milk and digests >90% in the presence of pepsin, as no detectable band is visible compared to the 0.02 mg/mL, 10% protein control (Figure 2A, B). Furthermore, the gel images show that the rhLF is highly pure (>99%) and no visible detection of host proteins is observed (Figure 2B). Bovine LF shows a similar trend (Supplemental Figure 1C). β-lactoglobulin, a known allergen previously shown to not digest in the presence of pepsin,^50^ appears completely intact, indicating it was not digested in these studies (Supplemental Figure 1D), confirming accuracy of the assay. We further verified digestion of the proteins using an automated western blot technique (Jess™, ProteinSimple, Bio-Techne). Importantly, this test further confirmed that Helaina rhLF digests in a comparable manner to hmLF with complete digestion of intact protein observed within 20 minutes of digestion (Figure 2C, D) and significant overlap of digestion products indicating similar peptide identities between rhLF and hmLF (Figure 2D). These data also corroborate the high purity of the rhLF and a lack of host protein presence. Interestingly, one peptide survived pepsin digestion with an approximate molecular weight of 2-3 kDa (Figure 2D) which correlates with published literature as lactoferricin,^51^ and remains intact throughout the entire 60-minute exposure (Supplemental Figure 2E). Additionally, quantification of intact proteins shows similar amounts between rhLF and hmLF throughout the digestion procedure with no intact protein remaining by 10 minutes for rhLF and 20 minutes for hmLF (Figure 2E). A pepsin-only control (no LF) was used to distinguish LF-specific peaks in the chromatogram from non-specific peaks (Supplemental figure 1F). Here, pepsin can be detected and is shown at ∼41 kDa as well as an unknown artifact at ∼27 kDa (Supplemental figure 1F). Finally, to further corroborate these findings, a standard western blot analysis was performed and showed similar results (Supplemental Figure 1G). Overall, these data indicate that Helaina rhLF digests comparably to hmLF, both showing low allergenic risk potential, and further suggests similar digestibility and potential bioavailability between the proteins.

**Figure 2.**
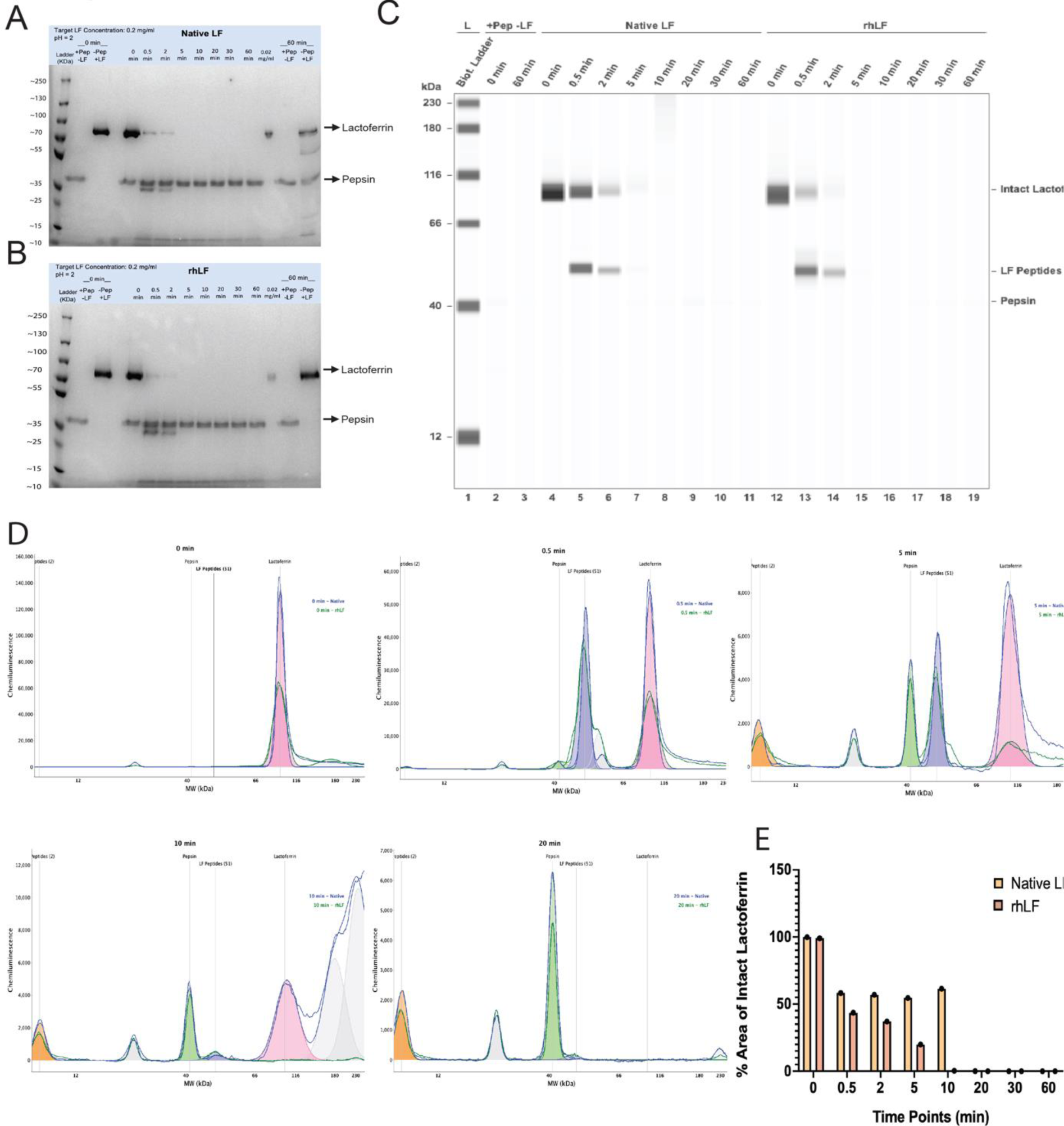
Helaina rhLF Digests in a Comparable Manner to Native Human LF and Shows Low Allergenic Risk Potential in the Pepsin Digestion Model. **A-B**: Coomassie Brilliant Blue stained SDS-PAGE gel showing the digestion of hmLF (A) or rhLF (B) in SGF at 10 units of pepsin per μg test protein at pH 2.0. Lane 1, protein ladder; Lane 2, pepsin control (no LF) at 0 minutes; Lane 3, protein control without pepsin at 0 minutes; Lane 4, empty; Lanes 5-12, digestion at 0, 0.5, 2, 5, 10, 20, 30, and 60 mins; Lane 13, protein at 10% concentration (0.02 mg/mL); Lane 14, pepsin only control (no LF) at 60 minutes; Lane 15, protein control without pepsin at 60 minutes. **C**. Jess™ gel blot showing digestion results for native hmLF and rhLF. Lane 1, protein ladder; Lane 2, pepsin control (no LF) at 0 minutes; Lane 3, pepsin control (no LF) at 60 minutes; Lanes 4-11, hmLF digestion at 0, 0.5, 2, 5, 10, 20, 30, and 60 mins; Lanes 12-19, rhLF digestion at 0, 0.5, 2, 5, 10, 20, 30, and 60 mins. **D-E**. Jess™ histogram of rhLF and hmLF following digestion at 0, 0.5, 5, 10, and 20 mins. **F**. Quantification of intact proteins of hmLF and rhLF following digestion at 0, 0.5, 2, 5, 10, 20, 30, and 60 mins from the Jess™ method.

## DISCUSSION

Novel foods require thorough safety evaluation prior to their introduction into the food supply, and the Codex Alimentarius Commission’s weight-of-evidence approach to evaluate allergenicity risk potential provides a framework by which to evaluate this aspect of the safety profile of novel proteins.^30,31^ To this end, we conducted a rigorous literature search, several bioinformatics exercises using AllergenOnline and BLASTP, and an in vitro digestion study using simulated gastric conditions to evaluate the potential allergenic risk of Helaina rhLF including its residual proteins expressed in *K. phaffii*. Neither the literature search, nor the bioinformatics exercises and pepsin digestion study suggest Helaina rhLF to pose allergenic risk to consumers, based on allergens known in 2023.

Literature searches for the primary protein of interest, human lactoferrin, did not identify studies showing allergy to this protein, even though the amino acid sequence is moderately identical to known potential allergens, bovine lactoferrin and ovo-transferrin. For example, there are no publications showing IgE cross-reactivity to hLF from sera of those allergic to bovine milk. But, in rare cases, bLF is the target of IgE binding from those allergic to cow’s milk outright,^52^ or due to α-gal carbohydrate on the asparagine linked sites on bLF.^53^ The other suggested cross-reactive allergen based on bioinformatics searches is ovo-transferrin, or Gal d 3, a major allergen naturally found in egg-white that binds IgE.^54,55^ Importantly, a study by Zhou, et al evaluated potential serum IgE binding between human LF (rhLF from transgenic cows or hLF) or bovine LF and sera from individuals allergic to milk or egg and found no visible cross-reactivity of rhLF, hLF, or bLF to the test serum samples^34^. The paper that suggested a possible risk from human lactoferrin was for protein expressed in rice that has cross-reactive carbohydrate determinants on some asparagine-linked glycosylation sites.^42^ Interestingly, the rice expressed protein did not elicit basophil activity and there was no proof of allergy to that product. Literature searches also did not identify studies of clinical allergy to the host source of *Komagataella phaffii* or using the previous name, *Pichia pastoris*.

Asparagine-linked glycans in eukaryotes can influence laboratory results in evaluating potential risks of allergy, such as the introduction of CCD in rice-produced hLF as described above.^42^ In the case of α-gal, which can occur in mammalian hosts, studies have shown this allergenic glycan in bovine milk, γ-globulin, lactoferrin, and lactoperoxidase.^53^ In the study evaluating rhLF produced in transgenic cows^34^ which would likely contain α-gal, no increased allergenic risk was observed, including no specific IgE binding in serum. Helaina-evaluated glycans detect neither CCD nor α-gal and contain only N-linked glycans that are oligomannose. While some studies have linked mannose binding the mannose receptor on DC to increased expression of Th2 cytokines, no definitive allergic testing was performed or identified.^56,57^ Further, Kreer et al. did a thorough analysis of N-glycans from *K. phaffii* and found no proof for mannose receptor on DC triggering allergic reactions.^44^ Therefore, Helaina rhLF glycans are unlikely to pose a significant risk of actual (α-gal) or perceived (CCD, mannose) allergen epitopes.

The current study identified that Helaina rhLF is a very pure protein with ≥99% of the protein content being the target protein, hLF, which is not linked with allergy. Due to low abundance of host proteins (<1% total) and the most abundant protein being only 0.045% of the total protein content, allergic responses against host proteins is unlikely. Nevertheless, a bioinformatics approach was utilized to determine allergenic potential of *K. phaffii* host proteins. The Codex Alimentarius criteria to consider potential risks of IgE cross-reactivity and clinical co-reactivity is 35% identity over 80 amino acids. As presented by Abdelmoteleb et al., these criteria are too low for evaluating potential risks of novel foods.^58^ There are many proteins conserved in evolution, and allergy across species is much more restricted to proteins that are not loose homologues. For most protein types, sequences of much greater than 50% identity is needed over most of the length of proteins for probable cross-reactivity.^58^ This risk is further reduced with AA sequence identity linked to human proteins, as these are likely to induce tolerance. These were the criteria considered when evaluating cross-reactivity and allergenic risk for the *K. phaffii* proteins identified in Helaina rhLF. The BLASTP matches of the host proteins to NCBI Protein did not uncover any matches to proteins that are likely allergens, and that are not in the AllergenOnline version 22 database. There were no high identity matches to allergens that would suggest potential clinical relevance or risks based on either the AllergenOnline sequence alignments or the NCBI Protein BLASTP searches. There are some interesting alignments to evolutionarily conserved proteins, yet those are unlikely to be allergens as human consumers would react to a wide variety of food sources that contain these proteins, which is rarely, if ever, documented.

Several studies have demonstrated pepsin resistance in food allergens, therefore showing a correlation between allergenic potential and resistance to pepsin digestion.^37,38,59^ For example, these studies have shown that allergenic proteins, such as β-lactoglobulin, are resistant to pepsin digestion; whereas non-allergenic proteins, such as horseradish peroxidase, concanavalin A, and soybean leghemoglobin, are susceptible to pepsin digestion. Previous work by Almond et al.,^60^ and Ofori-Anti, et al.,^38^ showed that rhLF (expressed in rice) and/or human LF are readily digested in the presence of pepsin. In our current studies, pepsin digestion modeling showed that Helaina rhLF was also rapidly digested in simulated gastric fluid. Importantly, its digestibility profile was comparable to hmLF, thus demonstrating a low allergenic risk. Interestingly, digested products (Helaina rhLF and hmLF) contained a small peptide that survived the entire digestion procedure, which corresponds in molecular weight to lactoferricin, a known peptide of lactoferrin.^51^

Although the existing evidence suggests a low allergenic risk potential of Helaina rhLF, Effera™, it cannot prove a nonexistent allergy risk. Nonetheless, these results do not support the need for further clinical testing or serum IgE binding to evaluate Helaina rhLF for risk of food allergy.

