## Supplemental Figure 1 for "Evaluation of the potential food allergy risks of recombinant human lactoferrin expressed in *Komagataella phaffii*"

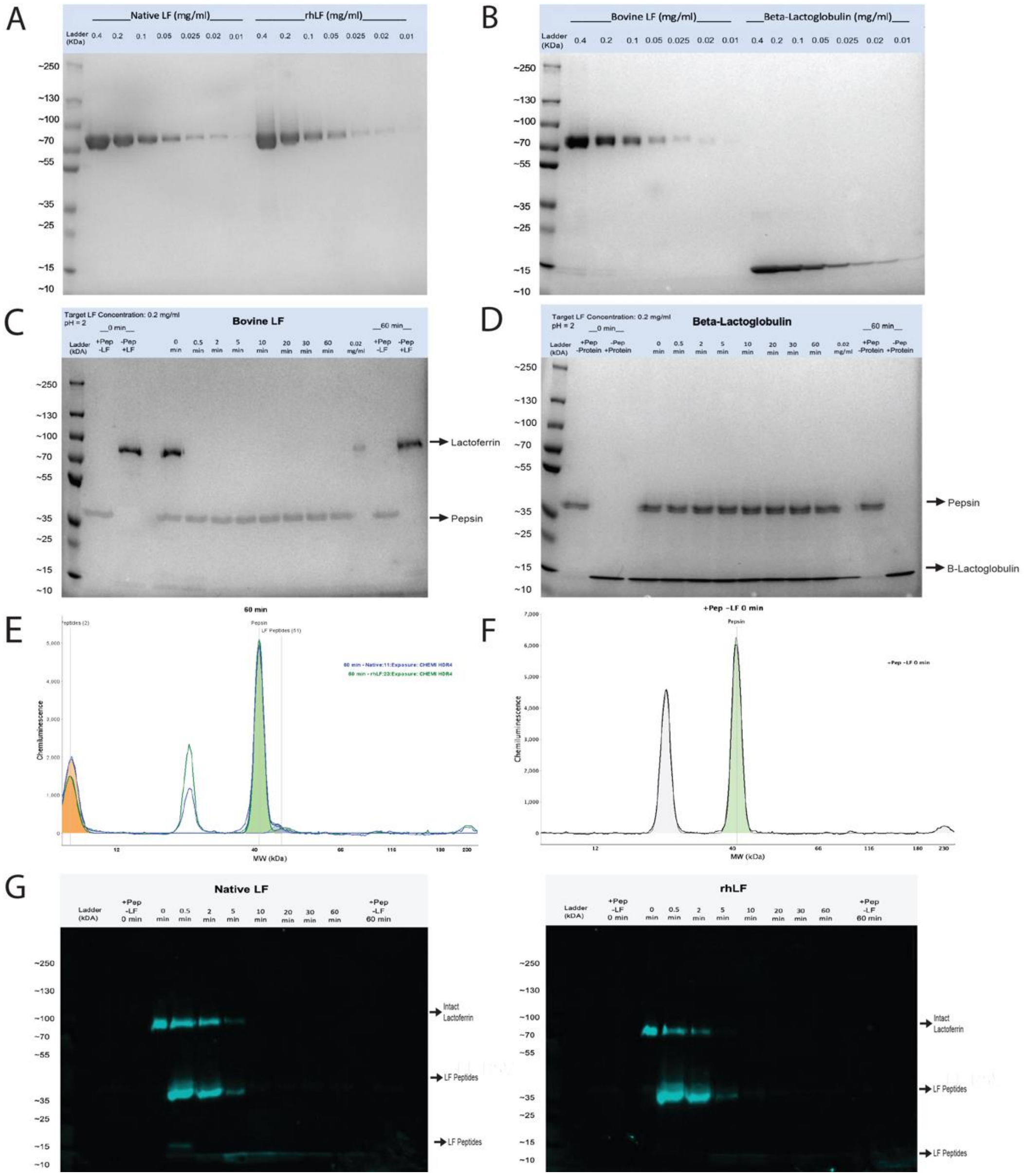


**Supplemental Figure 1**. **A-B**. Coomassie Brilliant Blue stained SDS-PAGE gel showing the serial dilution of target proteins from 200% to 5% of total protein. Proteins were loaded 10 µL per lane. Lane 1, protein ladder; Lane 2,9, 200% of total protein (4 µg); Lane 3,10, 100% of total protein (2 µg); Lane 4,11, 50% of total protein (1 µg); Lane 5,12, 25% of total protein (0.5 µg); Lane 6,13, 12.5% of total protein (0.25 µg); Lane 7,14, 10% of total protein (0.125 µg); Lane 8,15, 5% of total protein (0.0625 µg). A. Native LF (left) and rhLF produced from *Komagataella Phaffii* (right). B. Bovine LF (left) and beta-lactoglobulin (right). **C-D**. Coomassie Brilliant Blue stained SDS-PAGE gel showing the digestion of bovine LF (C) or beta-lactoglobulin (D) in SGF at 10 units of pepsin activity per microgram test protein at pH 2.0. Lane 1, protein ladder; Lane 2, pepsin control (no LF) at 0 minutes; Lane 3, protein control without pepsin at 0 minutes; Lane 4, empty; Lanes 5-12, digestion at 0, 0.5, 2, 5, 10, 20, 30, and 60 mins; Lane 13, protein at 10% concentration (0.02 mg/mL); Lane 14, pepsin only control (no LF) at 60 minutes; Lane 15, protein control without pepsin at 60 minutes. **E**. Jess™ histogram of pepsin alone (no LF) at 60 minutes. **F**. Fluorescent Western blot showing the digestion of native hmLF and rhLF in SGF at 10 units pepsin per microgram test protein at pH 2.0. Protein was loaded 10 µL per lane as pre- or post-digestion concentrate. Lane 1, protein ladder; Lane 2, empty; Lane 3, pepsin control (no LF) at 0 minutes; Lane 4, empty; Lanes 5-12, digestion at 0, 0.5, 2, 5, 10, 20, 30, and 60 mins; Lane 13, empty; Lane 14, pepsin only control (no LF) at 60 minutes.
